# MAE-UNETR++: Masked Autoencoder Pretraining for 3-D Lung Nodule Segmentation

**DOI:** 10.64898/2026.06.17.733000

**Authors:** Vinayak Savant, Yue Wang, Jianhua Xuan

## Abstract

Voxel-level annotation for volumetric medical imaging is expensive and difficult to scale, which makes training highcapacity 3-D segmentation models challenging in practice. Transfer learning (TL) from large public datasets is a common remedy, but it can under-perform when the source domain differs from the target anatomy and acquisition characteristics, as is often the case for pulmonary nodules. In this work, we propose a masked autoencoder (MAE) pretraining-based approach to break the data efficiency wall of domain difference and present a focused empirical study of domain-specific self-supervised learning (SSL) for 3-D lung nodule segmentation. We evaluate two experimental settings: first, Masked Autoencoder (MAE) pretraining versus random initialization across representative baselines; second, MAE versus Decathlon TL for UNETR++ while testing whether MAE-based pretraining also benefits a CNN baseline (V-Net). MAE pretraining on target-domain CT volumes achieves a Dice Similarity Coefficient (DSC) of 0.307, outperforming random initialization (0.136) and Decathlon weights (0.257). In addition, MAE improves the stability of V-Net in a “low-data” regime (i.e., with “insufficiently labeled” data), increasing DSC from 0.010 to0.071. Overall, these results suggest that MAE-based pretraining can provide a practical and robust initialization strategy for volumetric segmentation when labeled data are limited.

## I. Introduction

Lung cancer remains a leading cause of cancer-related mortality worldwide, and early detection is closely linked to improved outcomes [1], [2]. Low-dose CT screening has, therefore, increased interest in automated analysis pipelines, where accurate segmentation of pulmonary nodules supports measurement, follow-up, and clinical decision-making. Despite strong progress in 3-D deep learning, nodule segmentation remains a challenging setting for data-efficient training, nodules are small relative to the volume, boundaries can be ambiguous, and voxel-level labeling is time-consuming and expensive.

This problem is amplified as model architectures grow more expressive. Vision Transformers (ViTs) can model global context effectively, but stable representation learning typically benefits from large-scale training [3]. Conversely, CNNs embed locality and translation equivariance, which are often advantageous in limited-data settings. Yet, even well-established volumetric CNNs such as V-Net [7] can struggle when the positive class occupies a tiny fraction of the crop, making optimization sensitive to initialization and early training dynamics.

A widely used workaround is transfer learning (TL); for example, initializing from weights trained on the Medical Segmentation Decathlon (MSD) [20]. However, Decathlon pretraining spans heterogeneous anatomies, lesion scales, and acquisition protocols. For pulmonary nodules, small structures with diverse morphology e.g., juxta-pleural and spiculate patterns and inter-reader variability, this mismatch can limit both convergence behavior and final segmentation performance.

In this paper, we propose to use Masked Autoencoder (MAE) pretraining [5] as a domain-specific alternative and evaluate its effectiveness for lung nodule segmentation. MAE learns representations by reconstructing masked regions of an input volume using unlabeled data. Reconstructing missing anatomy encourages the encoder to internalize lung-specific structures and intensity statistics, which are precisely the priors that are hardest to learn from scarce labels. This paper makes three practical contributions: we provide a controlled evaluation of MAE pretraining for 3-D lung nodule segmentation using UNETR++ [4] on the LIDC-IDRI dataset [14]; we benchmark MAE against random initialization and Decathlon transfer learning [20]; and we test whether masking-based priors generalize beyond Transformers by applying MAE-based pretraining to a CNN baseline (V-Net) [7], focusing on convergence and stability in a low-data regime. Consistent with this scope, we study MAE primarily as an initialization strategy rather than as an architectural contribution, and focus on whether target-domain self-supervision improves optimization and segmentation quality under limited labels.

## II. Related Work

U-Net [6] and its volumetric extensions popularized encoder decoder segmentation with skip connections for biomedical imaging. V-Net [7] introduced residual-style volumetric feature learning and remains a strong CNN baseline, in part because convolutional inductive bias can be beneficial when data are limited. More recently, Transformer-based approaches have been adapted for 3-D medical segmentation to capture global context, including UNETR [9]. UNETR++ [4] improves efficiency and accuracy through Efficient Paired Attention (EPA) and multi-scale feature modeling, making it an appealing backbone when both global context and practical computation are important.

Self-supervised learning (SSL) is increasingly being used to reduce the reliance on labels. Contrastive approaches [10], [11] and masked modeling [5], [12] learn transferable representations without manual annotation. Masked modeling is particularly well-suited for medical volumes: reconstruction encourages anatomical consistency and intensity-aware priors rather than task-specific shortcuts. Pretraining strategies for transformer backbones in medical imaging, including Swin UNETR pretraining [13], more recent hierarchical masked pretraining for 3D medical images [21], [23], and recent reevaluations of MAE-style pretraining at larger scale [22], support the premise that SSL can improve data efficiency, but the relative benefit of domain-specific SSL versus supervised transfer remains task-dependent. In 3-D segmentation, Decathlon transfer learning [20] is widely used, yet its effectiveness depends on alignment between source tasks and the target distribution. For nodules, domain shift can arise from differences in lesion scale, reconstruction kernels, and preprocessing conventions, motivating a direct comparison between Decathlon initialization and MAE pretraining on unlabeled target-domain scans. We also note that UNETR++ is itself a recent 2024 Transformer backbone and is included here as a modern reference point. Our goal, however, is not to compare many architectures, but to isolate initialization under a fixed recent Transformer backbone and a representative volumetric CNN baseline. Broader comparisons to additional SSL methods and backbones remain important future work.

## III. Methods

Our primary backbone of the proposed approach is UN-ETR++ [4]. The encoder processes 3-D patch embeddings through hierarchical Transformer stages; Efficient Paired Attention (EPA) reduces computational overhead while maintaining global context, and the decoder combines multiscale representations via skip connections to produce high-resolution segmentation masks. We use V-Net [7] as a representative volumetric CNN baseline because its inductive bias is often favorable in limited-data settings, making it a strong reference point when assessing whether initialization and learned priors are bottlenecks under extreme class imbalance. Throughout this paper, V-Net (scratch) refers to random initialization without transfer learning or self-supervised pretraining, trained under the same supervised segmentation setup as the other baselines.

Fig. 1 provides a block diagram of the pipeline. We first use unlabeled target-domain cropped volumes to learn a strong initialization via self-supervision, and then fine-tune the same encoder within a segmentation model using the available voxel-level masks. This staging is motivated by the nodule setting itself: the foreground occupies a small portion of the cropped volume, so learning stable anatomical and intensity priors early can reduce the burden on scarce labeled supervision and improve downstream optimization.

**Fig. 1.**
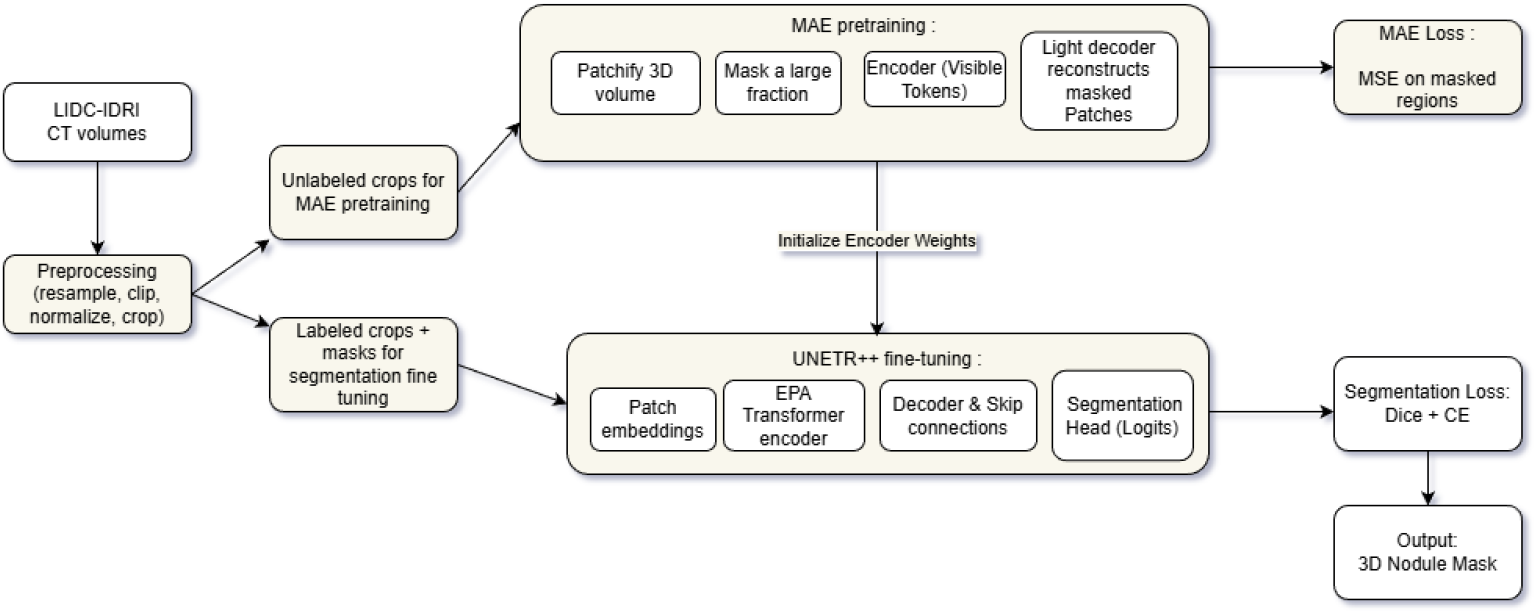
Overview of the proposed approach. MAE self-supervised pretraining learns lung-specific priors from unlabeled target-domain crops by reconstructing masked regions. The pretrained encoder weights are then used to initialize UNETR++ for supervised fine-tuning, producing a 3-D nodule mask optimized with a Dice+CE segmentation objective.

At a high level, the MAE stage is trained to reconstruct missing contents so the encoder is forced to encode contextual structure rather than relying on label-driven shortcuts. The resulting encoder weights are then used to initialize UNETR++ for supervised fine-tuning, where EPA-based attention and the decoder path with skip connections recover spatial detail and yield dense segmentation logits. This design keeps the method simple: the only change between stages is the learning objective and the availability of labels, while the encoder representation is carried forward.

For self-supervised pretraining, we use masking-based MAE [5] as shown in Fig. 1. The input volume *X* is partitioned into non-overlapping 3-D patches, and a large fraction of patches are masked. The encoder processes only visible patches, and a lightweight decoder reconstructs the masked content, encouraging lung-specific structural and texture priors from unlabeled scans. We use a high masking ratio (consistent with common MAE practice) and pre-train for 60 epochs on unlabeled target-domain crops. The reconstruction objective is the Mean Squared Error (MSE) over masked regions:

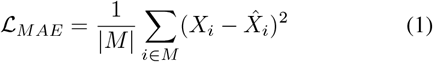

where *M* denotes masked elements and 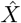 is the reconstruction.

For supervised segmentation, we fine-tune using voxel-level masks. Because nodules occupy a small fraction of each crop, we optimize a combined loss that balances overlap quality and voxel-wise stability as follows:

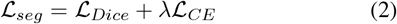

where *L*_CE_is the binary cross-entropy for binary masks and *λ* = 1.0. In our experiments, the segmentation fine-tuning is run for 150 epochs. The Dice loss function is defined by:

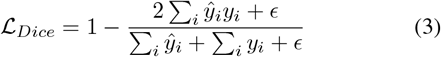

with *ϵ* for numerical stability.

## IV. Experimental Results

We benchmark on LIDC-IDRI [14], a thoracic CT dataset with multi-radiologist annotations. To standardize geometry across scanners and protocols, volumes are resampled to an isotropic resolution of 1.0 × 1.0 × 1.0 mm using cubic spline interpolation. LIDC scans can vary substantially in intensity scaling and stored ranges, so we apply a lung-relevant clipping step followed by an adaptive normalization procedure that accommodates both raw Hounsfield Unit (HU) volumes and scans that may already be normalized. This aims to preserve nodule-to-parenchyma contrast while producing a stable input distribution for training. For segmentation experiments, we extract fixed-size 3-D crops centered on the nodule region to preserve local context while keeping memory usage feasible, and we use 128 × 96 × 96 crops throughout unless noted otherwise.

All experiments are trained and evaluated on high-performance computing infrastructure. We use a patient-level split (80% training, 20% validation) to avoid leakage across subjects. Mixed-precision training (AMP) is enabled to improve throughput and maintain numerical stability, and we apply lightweight 3-D augmentations (axis flips, 90-degree rotations, and mild intensity perturbations) that preserve anatomy while reducing overfitting to scanner- or orientation-specific cues. We report the best validation DSC achieved during training using the same selection protocol across all methods, and qualitative overlays and heatmaps are generated from validation samples selected by a fixed rule to reduce presentation bias. In the submitted manuscript, this protocol did not include a separate held-out test set, as our goal was to keep the study focused on a controlled comparison of initialization strategies under the same low-label training and validation setting. Accordingly, all reported DSC values correspond to the validation protocol described above.

**TABLE I.** Experiment I: Impact Study of the Initialization or Pre-training Strategy.

| Model | Initialization | Dice Score ↑ |
| --- | --- | --- |
| V-Net | Random (Scratch) | 0.019 |
| UNETR++ | Random (Scratch) | 0.136 |
| **UNETR++ (Ours)** | **MAE (Pre-trained)** | **0.307** |

The first experimental setting evaluates whether MAE pretraining improves UNETR++ relative to training from scratch while also providing a reference CNN baseline. UNETR++ trained from random initialization reaches a DSC of 0.136, whereas MAE initialization increases performance to 0.307, indicating that representation learning priors substantially improve both convergence behavior and the final overlap in this low-label regime. V-Net trained from scratch performs poorly (0.019), which is consistent with the difficulty of optimizing stable volumetric features when positive voxels are extremely sparse and boundaries are subtle.

The second experimental setting benchmarks domain-specific MAE against supervised transfer learning and tests whether masking-based priors extend to a CNN. For UN-ETR++, the transfer learning baseline initializes from MSD Task06 Lung weights and achieves a DSC of 0.257, while MAE initialization achieves a DSC of 0.310. This gap suggests that representations learned directly from unlabeled target-domain CT volumes align more closely with lung nodule appearance than weights transferred from heterogeneous supervised tasks. For V-Net, MAE-style pre-training improves DSC from 0.010 to 0.071, indicating that masking-based priors are not limited to Transformer backbones and can stabilize early optimization for CNNs under the same class imbalance constraints.

To complement the adopted protocol, we also conducted a checkpoint-based held-out evaluation on the first fold using the same crop size and preprocessing path for both submitted UNETR++ checkpoints. Of the 225 cases tested, MAE achieved a mean Dice of 0.3591 compared with 0.3056 for Decathlon, corresponding to an improvement of 0.0535. A bootstrap analysis over cases gave a 95% confidence interval of [0.0340, 0.0742]. Across a threshold sweep, MAE remained higher across operating points, suggesting that the gain is not driven by a single threshold choice.

Beyond aggregate DSC, the qualitative overlays and attention-style heatmaps provide a useful sanity check on how the different initializations behave. In Fig. 3, MAE-initialized training rises earlier and stabilizes more smoothly, which is consistent with the encoder starting from features that already capture lung specific context rather than learning those priors from sparse foreground supervision. The slice-level overlays in Fig. 2 align with this trend, as MAE fine tuning tends to produce more coherent boundaries and fewer fragmented predictions, especially around subtle margins where intensity contrast is weak. Finally, the heatmaps in Fig. 5-6 suggest a shift in model focus: MAE initialization yields activations that concentrate around the nodule region, whereas the baselines more often exhibit diffuse or off-target responses. Taken together, these visuals support the interpretation that MAE primarily improves optimization by supplying a stronger anatomical prior early in training, which then translates into more reliable localization and delineation during fine tuning.

**Fig. 2.**
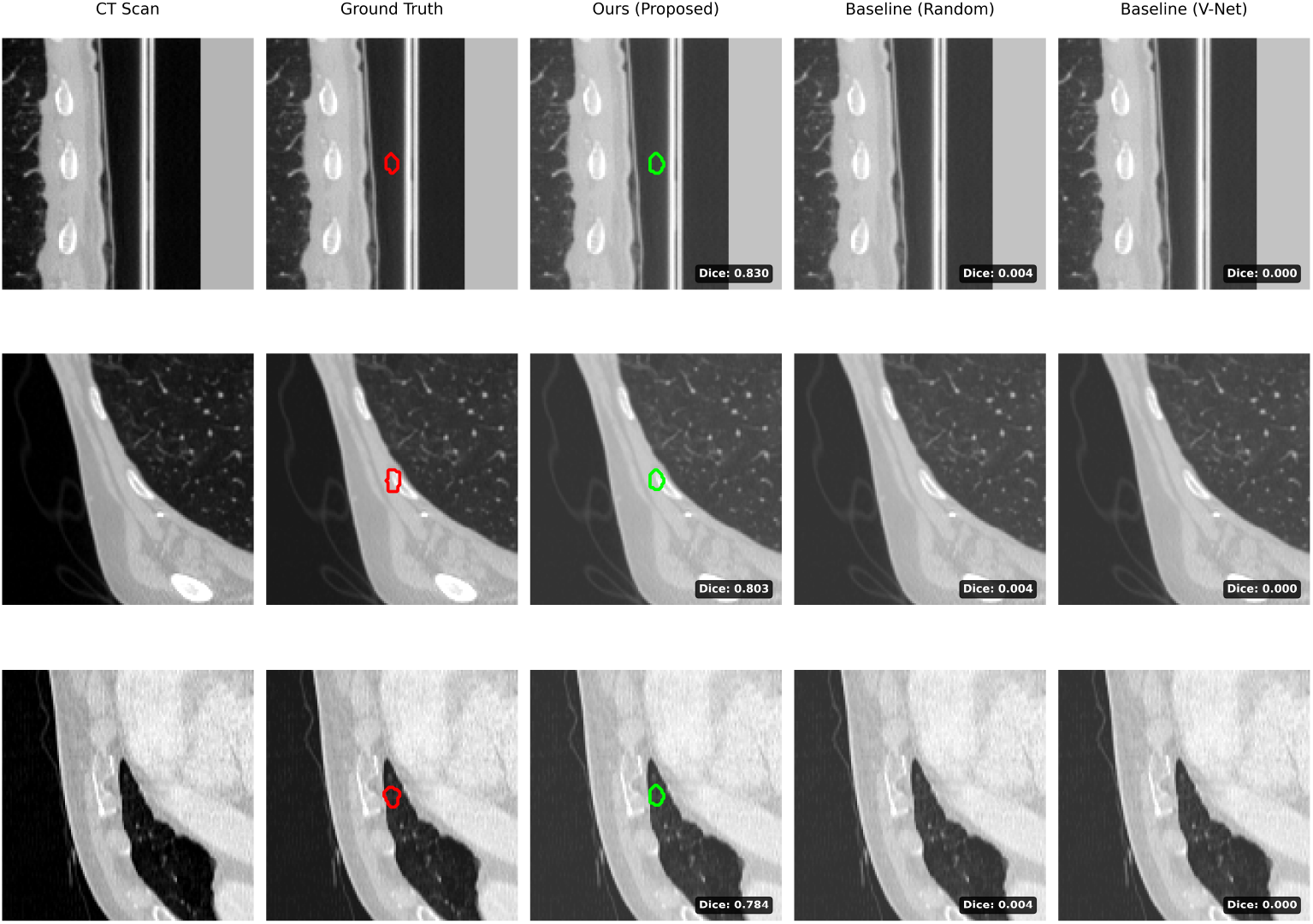
Qualitative Segmentation Results. (Left) CT slice and ground truth (Red). (Middle) UNETR++ MAE (Green) delineates the nodule boundary with strong overlap. (Right) Baselines under-segment or miss the region of interest, producing substantially lower overlap.

**Fig. 3.**
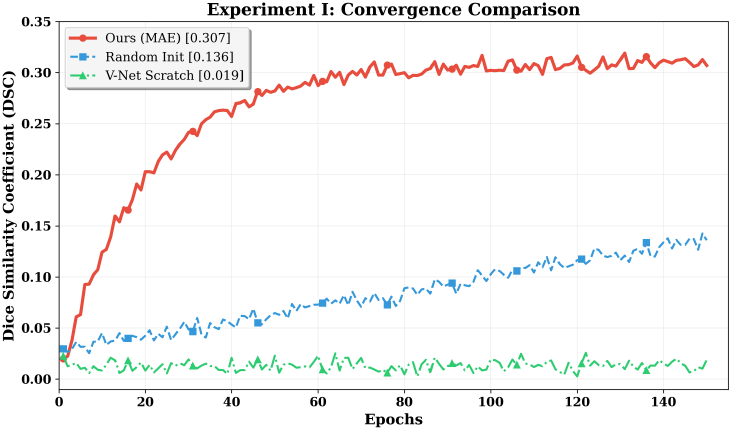
Convergence Comparison. Training dynamics comparing MAE initialization against baselines. MAE rises earlier and reaches a higher Dice plateau (0.307) than random initialization, indicating improved optimization and a better final solution.

**Fig. 4.**
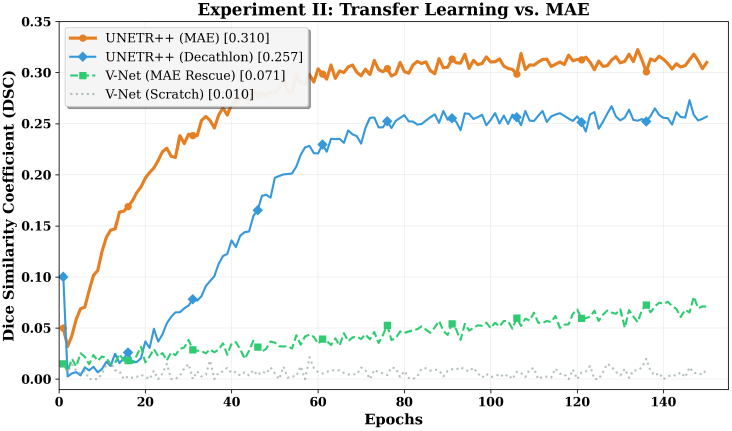
Experiment II Results. Domain-specific MAE pre-training outperforms Decathlon initialization for UNETR++. For V-Net, MAE improves training stability and raises DSC compared to scratch training.

**Fig. 5.**
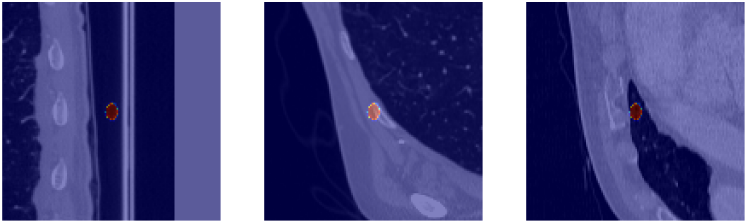
Model Focus (MAE). Heatmap visualization showing concentrated activation around the nodule region, consistent with localized decision-making aligned to the target structure.

**Fig. 6.**
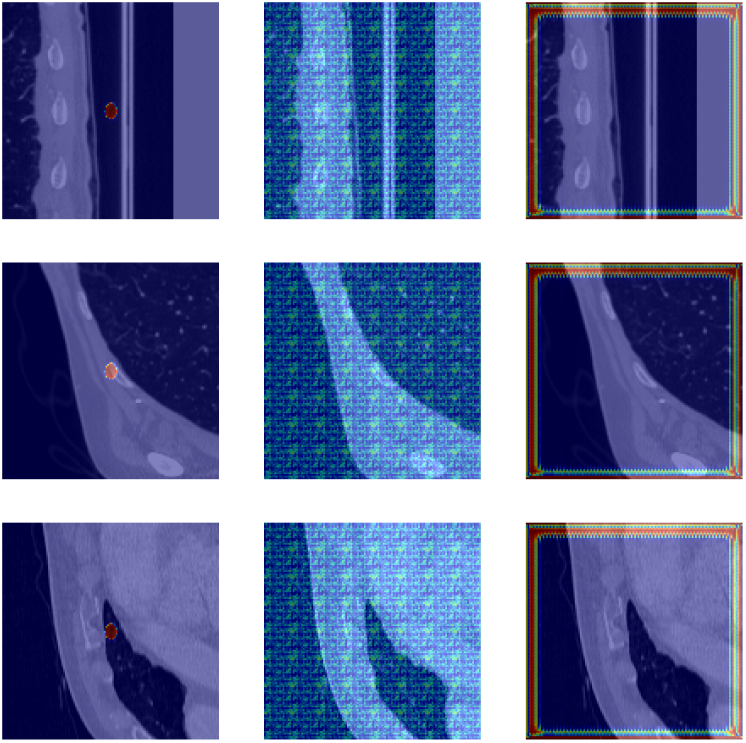
Comparative Heatmaps. Baselines often exhibit diffuse or off-target activation, whereas MAE initialization yields more localized and consistent focus on the region of interest.

**TABLE II.** Experiment ii: MAE VS. Decathlon & Architecture Generalization.

| Architecture | Method | Dice |
| --- | --- | --- |
| UNETR++ | Decathlon (TL, MSD Task06) | 0.257 |
|  | MAE (Ours) | **0.310** |
| V-Net | Scratch | 0.010 |
|  | MAE (Rescue) | **0.071** |

Related to false positives, the most common spurious pre-dictions were small leakage regions along adjacent vessels or pleural boundaries rather than diffuse off-target blobs. Qualitative inspection did not suggest that MAE increased this behavior relative to the baselines, which is consistent with the more localized activation patterns observed in the heatmap visualizations.

Figure 7 shows several representative difficult cases. The dominant failure modes are consistent across MAE and Decathlon: very small or low-contrast nodules, nodules adjacent to vessels or pleura where local intensities are confounded, and cases with higher annotation uncertainty where multiple plausible boundaries exist. In these cases, the models may under-segment, miss the lesion entirely, or leak into nearby structures.

**Fig. 7.**
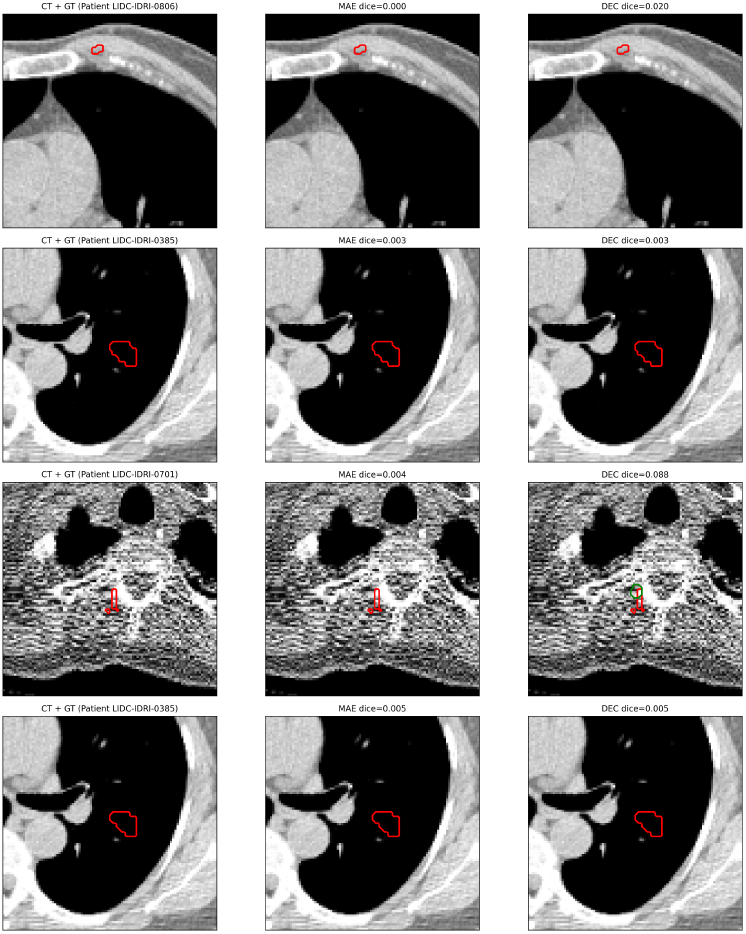
Representative difficult cases. Failure examples from the lowestperforming cases for MAE and Decathlon. Common failure modes include missed lesions and severe under-segmentation in very small or low-contrast nodules, nodules adjacent to vessels or pleura, and cases with boundary ambiguity.

These results suggest that the key bottleneck in this setting is not only architectural capacity but also the availability of lung-specific priors early in training. For UNETR++ [4], MAE more than doubles DSC relative to random initialization in the first setting and remains stronger than Decathlon transfer learning in the second setting. The V-Net [7] results reinforce this point from a different angle, indicating that masking-based pretraining can also stabilize optimization for a CNN baseline under severe class imbalance.

This study is intentionally empirical. The experiments are conducted on a single dataset using one recent Transformer backbone and one representative volumetric CNN baseline, and we do not compare against alternative SSL strategies such as SimCLR, DINO, or SparK. These choices were made to isolate initialization as the main variable, but broader architectural and dataset-level validation remains important future work.

## V. Conclusion

We have presented a targeted empirical evaluation of MAE pretraining for 3-D lung nodule segmentation on LIDC-IDRI using UNETR++ and V-Net as representative Transformer and CNN baselines. Across two experimental settings, domain-specific MAE initialization consistently improved DSC over random initialization and outperformed Decathlon transfer learning initialized from MSD Task06 (Lung) weights for UN-ETR++. In addition, MAE pretraining improved convergence for a CNN baseline (V-Net) in a low-data regime. We therefore view the contribution not as a new segmentation architecture, but as evidence that target-domain MAE pretraining can serve as a practical initialization strategy when voxel-level labels are scarce. Overall, the results support masking-based self-supervision on target-domain scans as a practical route to improved data efficiency and more stable optimization for volumetric medical segmentation.

For future work, we will further evaluate the robustness of the approach beyond DSC by reporting boundary-sensitive metrics such as surface distance and to test generalization under acquisition shifts and external cohorts. We also plan to scale training and pretraining to a larger dataset to better quantify how unlabeled data volume impacts downstream gains under fixed compute budgets. Beyond segmentation, we intend to extend the pipeline toward clinical characterization by adding a nodule classification module to predict benign versus malignant status, using segmentation-derived regions of interest and complementary radiographic cues. Finally, we will study sensitivity to masking schedules and pretraining duration to better map the trade-off between pretraining cost and downstream performance for both Transformer and CNN backbones.

## Acknowledgment

This work was supported by the National Institutes of Health (NIH) under Grants MH110504 and CA271891 to Y.W.

## References

[1] H. Sung et al., “Global cancer statistics 2020: GLOBOCAN estimates of incidence and mortality worldwide for 36 cancers in 185 countries,” CA: A Cancer Journal for Clinicians, vol. 71, no. 3, pp. 209–249, 2021.

[2] National Lung Screening Trial Research Team, “Reduced lung-cancer mortality with low-dose computed tomographic screening,” NEJM, vol. 365, no. 5, pp. 395–409, 2011.

[3] A. Dosovitskiy et al., “An image is worth 16x16 words: Transformers for image recognition at scale,” ICLR, 2021.

[4] A. Shaker et al., “UNETR++: Delving Into Efficient and Accurate 3D Medical Image Segmentation,” IEEE Transactions on Medical Imaging, vol. 43, no. 9, pp. 3377–3390, Sep. 2024, doi: 10.1109/TMI.2024.3398728.

[5] K. He et al., “Masked autoencoders are scalable vision learners,” CVPR, pp. 16000–16009, 2022.

[6] O. Ronneberger et al., “U-net: Convolutional networks for biomedical image segmentation,” MICCAI, pp. 234–241, 2015.

[7] F. Milletari, N. Navab, and S.-A. Ahmadi, “V-Net: Fully Convolutional Neural Networks for Volumetric Medical Image Segmentation,” in Proc. 2016 Fourth International Conference on 3D Vision (3DV), 2016, pp. 565–571, doi: 10.1109/3DV.2016.79.

[8] F. Isensee et al., “nnU-Net: a self-configuring method for deep learning-based biomedical image segmentation,” Nature Methods, vol. 18, pp. 203–211, 2021.

[9] A. Hatamizadeh et al., “Unetr: Transformers for 3d medical image segmentation,” WACV, 2022.

[10] T. Chen et al., “A simple framework for contrastive learning of visual representations,” ICML, 2020.

[11] K. He et al., “Momentum contrast for unsupervised visual representation learning,” CVPR, 2020.

[12] H. Bao et al., “Beit: Bert pre-training of image transformers,” ICLR, 2022.

[13] Y. Tang et al., “Self-supervised pre-training of swin transformers for 3d medical image analysis,” CVPR, 2022.

[14] S. G. Armato III et al., “The lung image database consortium (LIDC) and image database resource initiative (IDRI),” Medical Physics, vol. 38, no. 2, 2011.

[15] F. Ciompi et al., “Towards automatic pulmonary nodule management in lung cancer screening with deep learning,” Scientific Reports, 2015.

[16] A. A. A. Setio et al., “Pulmonary nodule detection in CT images: false positive reduction using multi-view convolutional networks,” IEEE TMI, 2016.

[17] K. Hara et al., “Can spatiotemporal 3d cnns retrace the history of 2d cnns and imagenet?” CVPR, 2018.

[18] W. Zhu et al., “Deepem: Deep 3d convnets with em for clustering without supervised labels,” MICCAI, 2018.

[19] H. Tang et al., “Nodulenet: Decoupled false positive reduction for pulmonary nodule detection and segmentation,” MICCAI, 2019.

[20] A. Antonelli et al., “The Medical Segmentation Decathlon,” Nature Communications, vol. 13, art. no. 4128, 2022, doi: 10.1038/s41467-022-30695-9.

[21] J. Zhuang, L. Wu, Q. Wang, P. Fei, V. Vardhanabhuti, L. Luo, and H. Chen, “MiM: Mask in Mask Self-Supervised Pre-Training for 3D Medical Image Analysis,” arXiv preprint arXiv:2404.15580, 2024.

[22] T. Wald, C. Ulrich, S. Lukyanenko, A. Goncharov, A. Paderno, L. Maerkisch, P. F. Jaäger, and K. Maier-Hein, “Revisiting MAE Pre-Training for 3D Medical Image Segmentation,” in Proceedings of the IEEE/CVF Conference on Computer Vision and Pattern Recognition (CVPR), 2025.

[23] Y. Ding, J. Wang, and H. Lyu, “Intensity-Spatial Dual Masked Autoencoder for Multi-Scale Feature Learning in Chest CT Segmentation,” arXiv preprint arXiv:2411.13198, 2024.

